# Paternal Metabolic Reversal Remodels Sperm RNA Profiles and Ameliorates Intergenerational Metabolic Disorder in Mice

**DOI:** 10.64898/2026.07.31.742153

**Authors:** Sheng Chen, Rubens Daniel Miserani Magalhaes, Zhuqing Wang, Fritz Cayabyab, Jinhyuk Choi, Eiji Yoshihara, Rui Wang, Hayden McSwiggin, Laura Chavez, Harry B. Rossiter, Rachelle Bross, Yanhe Lue, Christina Wang, Ronald S. Swerdloff, John R. McCarrey, Huili Zheng, Wei Yan

## Abstract

Paternal obesity increases metabolic risk in offspring, but whether this risk can be reduced by restoring paternal health before conception remains unresolved. We developed a within-sire induction-and-reversal model in outbred CD1 mice in which high-fat diet (HFD)-exposed males generated offspring before and after transition to an ingredient-matched control diet with voluntary exercise. HFD caused obesity, glucose intolerance, insulin resistance, and extensive remodeling of sperm mRNA, lncRNA, and sncRNA profiles, together with transcriptomic changes in metabolic tissues. Diet and exercise reversal normalized paternal metabolic indices and broadly restored tissue RNA profiles, although sperm retained a limited transcriptional memory of prior HFD exposure. Offspring sired before reversal developed sex-dependent metabolic dysfunction despite control-diet rearing, whereas offspring sired after reversal showed substantial improvement. These findings show that paternal metabolic risk is modifiable before conception and that this reversibility is linked to remodeling of sperm RNA. (**140 words**)

**Highlights:**

- Paternal HFD-Ex induces obesity, glucose intolerance and insulin resistance in CD1 males
- Sperm shows much stronger RNA response than four metabolic organs profiled
- Diet and exercise reversal restores metabolism and RNA profiles in sperm and four metabolic organs analyzed
- Offspring metabolic risk is reduced when sires conceive after reversal through diet and exercise intervention

**eTOC Blurb:** Chen, Magalhaes, et al. show that paternal metabolic recovery before conception remodels sperm RNA and reduces transmission of HFD-associated metabolic risk to offspring in a within-sire mouse model.

## Introduction

The developmental origins of health and disease (DOHaD) framework has traditionally emphasized the maternal gestational environment. Increasing evidence indicates that the father’s preconception metabolic state also contributes to offspring metabolic programming. Paternal obesity, high-fat diet, and metabolic dysfunction have been linked to altered adiposity, glucose tolerance, insulin sensitivity, and lipid homeostasis in offspring in rodent models.^1–5^ Human data are consistent with this view. In ART-conceived children, paternal BMI has been associated with offspring BMI, systolic blood pressure, and insulin resistance,^6^ and parent-offspring BMI associations at comparable developmental stages are not readily explained by gestational exposure alone.^7^ Nevertheless, preconception care remains focused primarily on maternal health, with limited attention to paternal metabolic status.^8,9^

Sperm carry epigenetic information in addition to the paternal genome. Mature sperm contain mRNAs, large noncoding RNAs (lncRNAs), microRNAs (miRNAs), tRNA-derived small RNAs (tsRNAs), rRNA-derived small RNAs (rsRNAs), Piwi-interacting RNAs (piRNAs), and other small noncoding RNAs (sncRNAs) that are delivered to the oocyte at fertilization.^10–16^ Functional studies support a role for sperm RNA in intergenerational inheritance. Sperm RNA from diet-challenged or stressed males can recapitulate aspects of offspring metabolic or behavioral phenotypes after zygote injection,^17–19^ and loss of the RNA methyltransferase DNMT2 disrupts sperm sncRNA-mediated transmission of HFD-induced metabolic traits.^20^ These findings identify sperm RNA as a plausible carrier of environmentally responsive paternal information.^21^

Where paternal information enters sperm remains an important mechanistic question. Two post-testicular models have been proposed: epididymal soma-to-spern RNA transfer through epididymosomes and mitochondrial DNA-dependent transcription in mature sperm. ^12,22,23^ However, both have been challenged by subsequent evidence.^24–26^ Caput epididymal sperm can support full-term development;^24,25^ many RNA differences observed during epididymal maturation can be explained by redistribution between sperm and cytoplasmic droplets;^27^ and mature sperm contain little to no intact mitochondrial DNA.^26^ ICSI using testicular sperm heads further indicates that diet-induced heritable metabolic information is already present before epididymal transit.^28^ These observations support a testicular encoding model in which the paternal metabolic state is translated into sperm-borne information during spermatogenesis.

A clinically central question remains unresolved: *whether an adverse paternal signal produced by metabolic disease can be reversed by improving paternal health before conception*. Prior studies have shown that paternal exercise or dietary intervention can ameliorate offspring metabolic abnormalities and alter sperm sncRNAs or chromatin marks.^29–31^ However, most studies compared independent paternal cohorts, leaving open the contribution of between-cohort variation. A stringent test requires the same male to generate offspring before and after recovery, so that the paternal state at conception can be isolated from genetic background and founder effects.

To address this question, we developed a within-sire induction-and-reversal model. Outbred CD1 males were exposed to HFD without exercise (HFD-Ex) to induce obesity and metabolic dysfunction, mated to generate offspring, then switched to an ingredient-matched control diet with voluntary exercise (ICD+Ex) and mated again after metabolic recovery (Figure 1A). We profiled both large and small RNAs in sperm and four metabolic tissues (islets, liver, muscle, and brain) across four paternal conditions: ICD+Ex, HFD-Ex, RICD (continued ICD+Ex), and RHFD (switching from HFD-Ex to ICD+Ex), and assessed metabolic phenotypes in F1 offspring reared on regular chow (Figure 1A). The design provides direct evidence that paternal metabolic risk is substantially reversible before conception and identifies sperm RNA remodeling as a molecular correlate of that reversibility.

**Figure 1.**
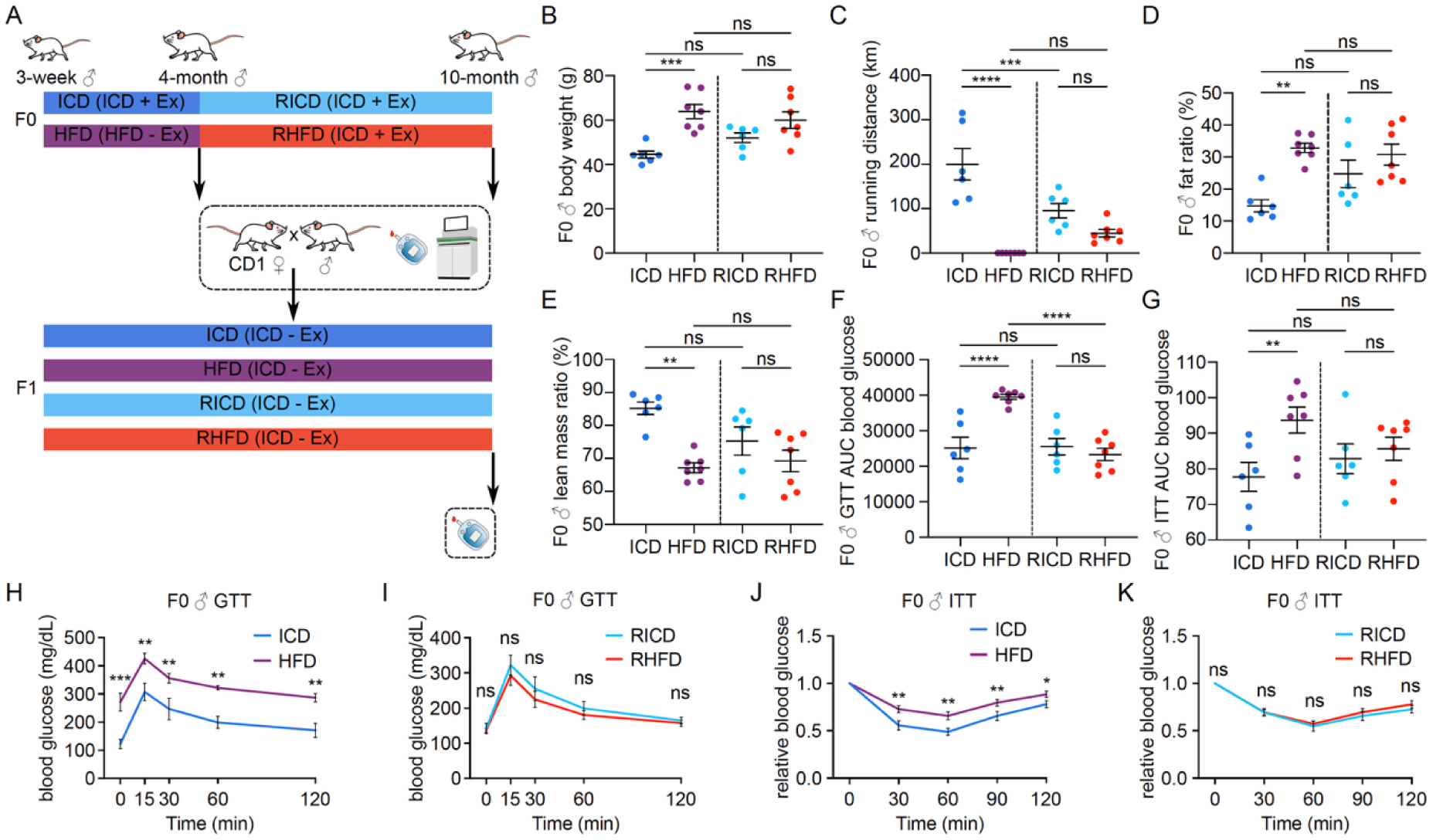
Experimental design and metabolic reversal in F0 males. (A) Schematic of the paternal induction-and-reversal paradigm. Three-week-old outbred CD1 males were acclimated for 1 week and randomized at approximately 4 weeks to ICD+Ex (ingredient-matched control diet with voluntary wheel running) or HFD-Ex (high-fat diet with locked wheel/no exercise) for 16 weeks. After phenotyping and first breeding, HFD-Ex males were transitioned to ICD+Ex (RHFD); age-matched controls remained on ICD+Ex (RICD) for at least 16 weeks before re-phenotyping and second breeding. All F1 offspring were maintained on regular chow without exercise. (B-E) Body weight (B), running distance (C), fat mass ratio (D), and lean mass ratio (E) in F0 males. (F and G) GTT AUC (F) and ITT AUC (G). (H-K) GTT curves before (H) and after (I) reversal and ITT curves before (J) and after (K) reversal. Data are mean ± SEM. *, **, ***, ****, and ns indicate p < 0.05, p < 0.01, p < 0.001, p < 0.0001, and not significant, respectively.

## Results

### Establishment of a paternal induction-and-reversal model in outbred CD1 mice

Outbred CD1 F0 males were purchased at 3 weeks of age, acclimated for 1 week, and randomized at approximately 4 weeks to either ICD+Ex, an ingredient-matched control diet with voluntary running (Envigo Teklad TD.120455; 16.8% kcal fat; 3.3 kcal/g), or HFD-Ex, a high-fat diet with a locked running wheel/no exercise (Envigo Teklad TD.08811; 44.6% kcal fat; 4.7 kcal/g). After 16 weeks, HFD-Ex males had increased body weight and fat mass ratio, reduced lean mass ratio, impaired glucose tolerance, and insulin resistance relative to ICD+Ex controls (Figures 1B-1H and 1J; Table S1).

After the first breeding, HFD-Ex males were switched to ICD+Ex for at least 16 weeks to generate the RHFD group, whereas age-matched ICD+Ex controls continued on the same regimen to generate the RICD group. Following reversal, RHFD males showed body composition, voluntary running, GTT responses, and ITT responses comparable to RICD controls (Figures 1B-1G, 1I, and 1K; Table S1). Thus, the model established matched paternal states before and after metabolic recovery and enabled comparison of offspring with the same paternal exposure history.

### HFD-Ex induces tissue-specific mRNA remodeling that is largely reversible

Bulk RNA-seq was performed on sperm, pancreatic islets, skeletal muscle, liver, and brain from ICD, HFD, RICD, and RHFD males. Principal-component analysis showed diet-and tissue-dependent separation of mRNA profiles (Figure 2A). Differential expression analysis identified HFD-responsive genes in all five tissues, with the strongest response in sperm (Figure 2B; Table S2). Comparison of HFD versus ICD with RHFD versus RICD showed extensive reversal in somatic tissues: all initial HFD-responsive mRNA changes were resolved in muscle, liver, and brain, and 94% were resolved in islets. Sperm showed partial but substantial recovery, with 79% of initial HFD-responsive mRNAs no longer differentially expressed after reversal (Figure 2B; Table S2). The 21% of sperm mRNAs that persisted after reversal were enriched for developmental, immune-response, and translational processes (Figure 2B; Table S3).

**Figure 2.**
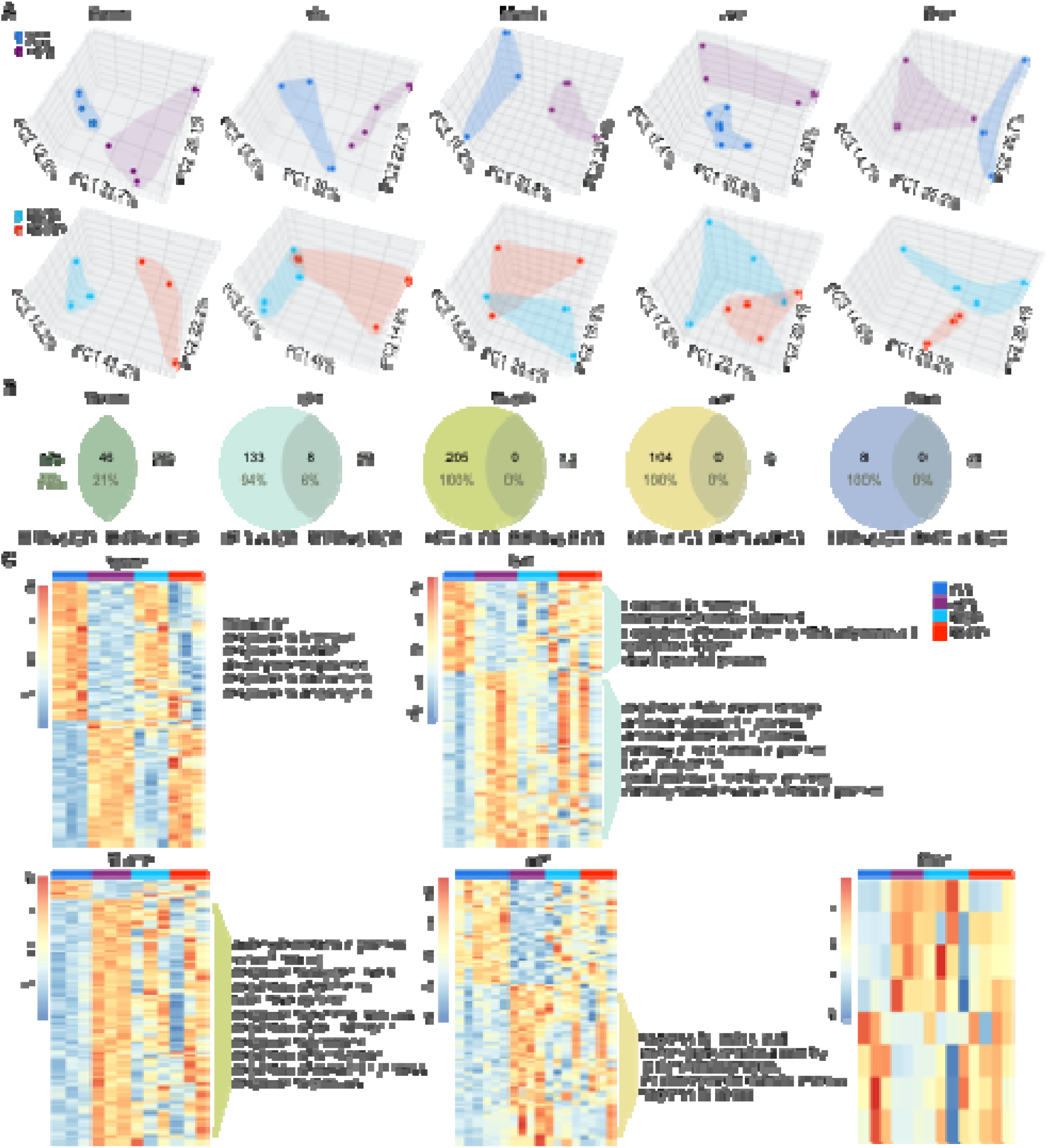
HFD-induced and reversal-associated mRNA profiles across F0 sperm and metabolic tissues. (A) PCA plots of mRNA profiles in sperm, pancreatic islets, skeletal muscle, liver, and brain from ICD, HFD, RICD, and RHFD males. (B) Venn diagrams showing differentially expressed mRNAs in HFD versus ICD and RHFD versus RICD comparisons. Initial HFD-responsive mRNAs were largely resolved after reversal, with recovery rates of 79% in sperm, 94% in islets, and 100% in muscle, liver, and brain. (C) Heatmaps and representative GO terms for differentially expressed mRNAs in each tissue. Differential expression was determined by DESeq2 with adjusted p < 0.05.

Although most HFD-associated changes were corrected, newly RHFD-versus-RICD differences were detected in sperm, islets, and brain (Figure 2B; Table S2). These differences occurred after systemic metabolic normalization and may represent residual transcriptional effects of prior exposure rather than persistent obesity. GO enrichment of HFD-responsive sperm mRNAs implicated translation, hormone response, cAMP signaling, calcium ion response, developmental processes, and stress-response pathways (Figure 2C; Table S3). Representative sperm genes with HFD-induced changes and reversal included *Slc14a2*, *Smarcd3*, and *Srek1* (Figure S1).

### LncRNA profiles show broad recovery with residual sperm-associated differences

LncRNA profiles displayed tissue-and condition-dependent separation similar to mRNAs (Figure 3A). Differential lncRNA expression was detected across all tissues, with high reversal rates following the diet and exercise intervention: approximately 95% in sperm and islets, 98% in muscle, and 100% in liver and brain (Figure 3B; Table S4). Heatmaps confirmed broad normalization of lncRNA profiles after reversal, while sperm retained a detectable subset of reversal-associated differences (Figure 3C). These data extend the reversible paternal RNA response beyond protein-coding transcripts and suggest that lncRNA content also reflects paternal metabolic state during spermatogenesis.

**Figure 3.**
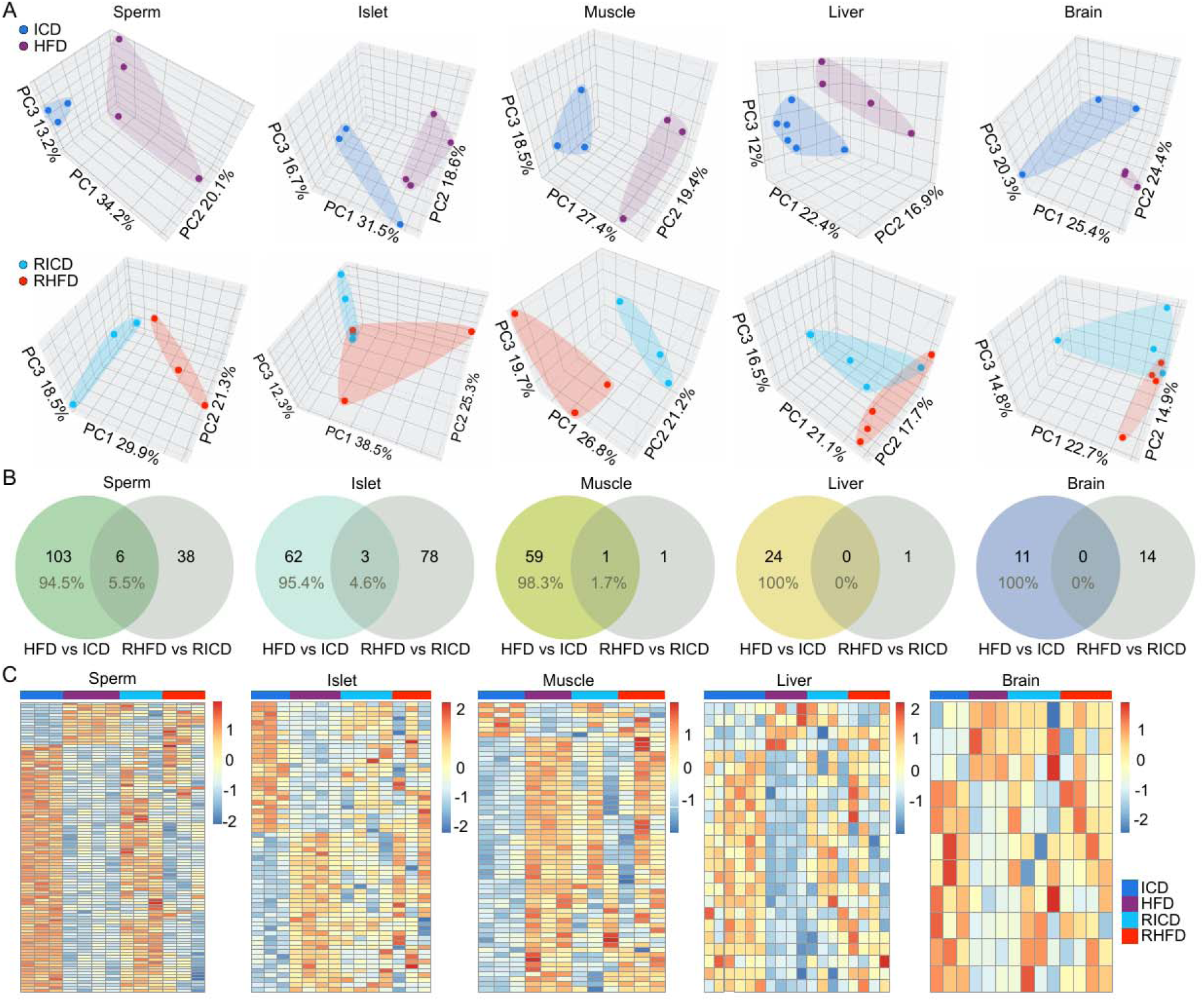
HFD-induced and reversal-associated lncRNA profiles across F0 sperm and metabolic tissues. (A) PCA plots of lncRNA profiles in sperm, pancreatic islets, skeletal muscle, liver, and brain. (B) Venn diagrams showing differentially expressed lncRNAs in HFD versus ICD and RHFD versus RICD comparisons. (C) Heatmaps of tissue-specific lncRNA responses across ICD, HFD, RICD, and RHFD groups. Differential expression was determined by DESeq2 with adjusted p < 0.05.

### HFD-Ex causes extensive sperm sncRNA remodeling that is largely reset by reversal

sncRNA-seq libraries were annotated with SPORTS1.1 and analyzed with edgeR. ^32^ Across the five tissues, sperm showed the greatest diversity of sncRNAs, dominated by piRNAs and rRNA-derived small RNAs (Figure 4A; Table S4). HFD-Ex produced a large shift in sperm sncRNA composition, with 144,999 differentially expressed sncRNA sequence features in HFD versus ICD sperm; these features were predominantly annotated as piRNAs and rsRNAs (Figures 4B and 4C; Table S5). By contrast, far fewer sncRNA changes were detected in muscle and islets, and no significant changes were detected in liver or brain under the same criteria (Figure 4C; Table S5).

**Figure 4.**
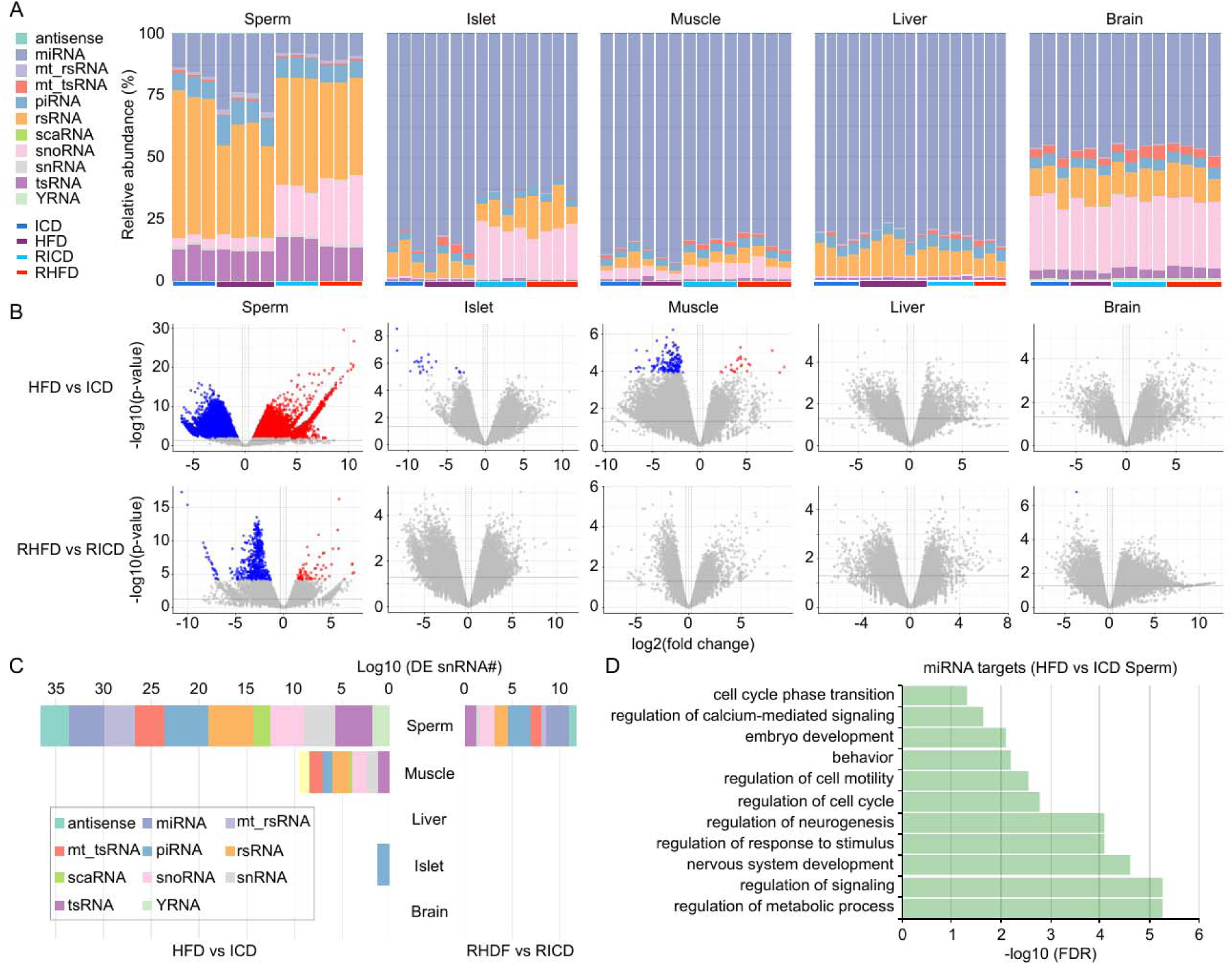
sncRNA remodeling by HFD-Ex and reversal. (A) Relative abundance of sncRNA subtypes across sperm, pancreatic islets, skeletal muscle, liver, and brain. (B) Volcano plots of differentially expressed sncRNAs in HFD versus ICD and RHFD versus RICD comparisons. (C) Summary of differentially expressed sncRNAs by tissue and biotype on a log10 scale. HFD-Ex altered 144,999 sncRNA sequences in sperm, whereas 869 remained differentially expressed in RHFD versus RICD sperm. (D) GO enrichment of predicted targets of HFD-responsive sperm miRNAs. Target genes were defined by overlap among miRWalk, TargetScan, and miRDB predictions. Differential expression was determined by edgeR with adjusted p < 0.05.

Diet and exercise reversal reduced the number of differentially expressed sperm sncRNA features, from 144,999 in HFD versus ICD sperm to 869 in RHFD versus RICD sperm (Figure 4C; Table S5). Correlation analysis further showed that HFD and ICD sperm sncRNA profiles were separated, whereas RHFD and RICD profiles were closely correlated (Figure S6). Thus, sperm sncRNAs are highly sensitive to paternal metabolic stress but are also strongly reset by restoration of a healthy metabolic state.

To assess potential functional consequences of HFD-responsive sperm miRNAs, we intersected predictions from miRWalk, TargetScan, and miRDB. This analysis linked 6,699 differentially expressed miRNA sequences to 560 high-confidence target genes (Table S6). GO enrichment implicated regulation of metabolic processes, signaling, neurogenesis, cell cycle, embryo development, and calcium-mediated signaling (Figure 4D; Table S6), consistent with pathways that could influence early development and later metabolic physiology.

### Paternal reversal reduces intergenerational metabolic transmission to F1 offspring

To determine whether the father’s metabolic state at conception affects offspring outcome, F0 males were timely mated with virgin CD1 females before and after reversal. All F1 offspring were maintained on regular chow without exercise from weaning, thereby separating offspring phenotype from direct HFD or exercise exposure.

At 10 months of age, male offspring from HFD sires had increased body weight and impaired glucose tolerance compared with male offspring from ICD sires (Figures 5A-5D). Male offspring from RHFD sires showed body weight and glucose tolerance that were substantially normalized toward RICD-derived controls (Figures 5A-5D; Table S1). Female offspring from HFD sires also displayed metabolic impairment, including glucose intolerance and altered insulin sensitivity (Figures 5E-5K; Table S1). After paternal reversal, female offspring showed improved GTT and ITT parameters, although the rescue was less complete than in males (Figures 5F, 5H, and 5I-5K; Table S1). These results indicate that paternal metabolic recovery before conception reduces intergenerational metabolic risk, with the extent of rescue differing between male and female offspring.

**Figure 5.**
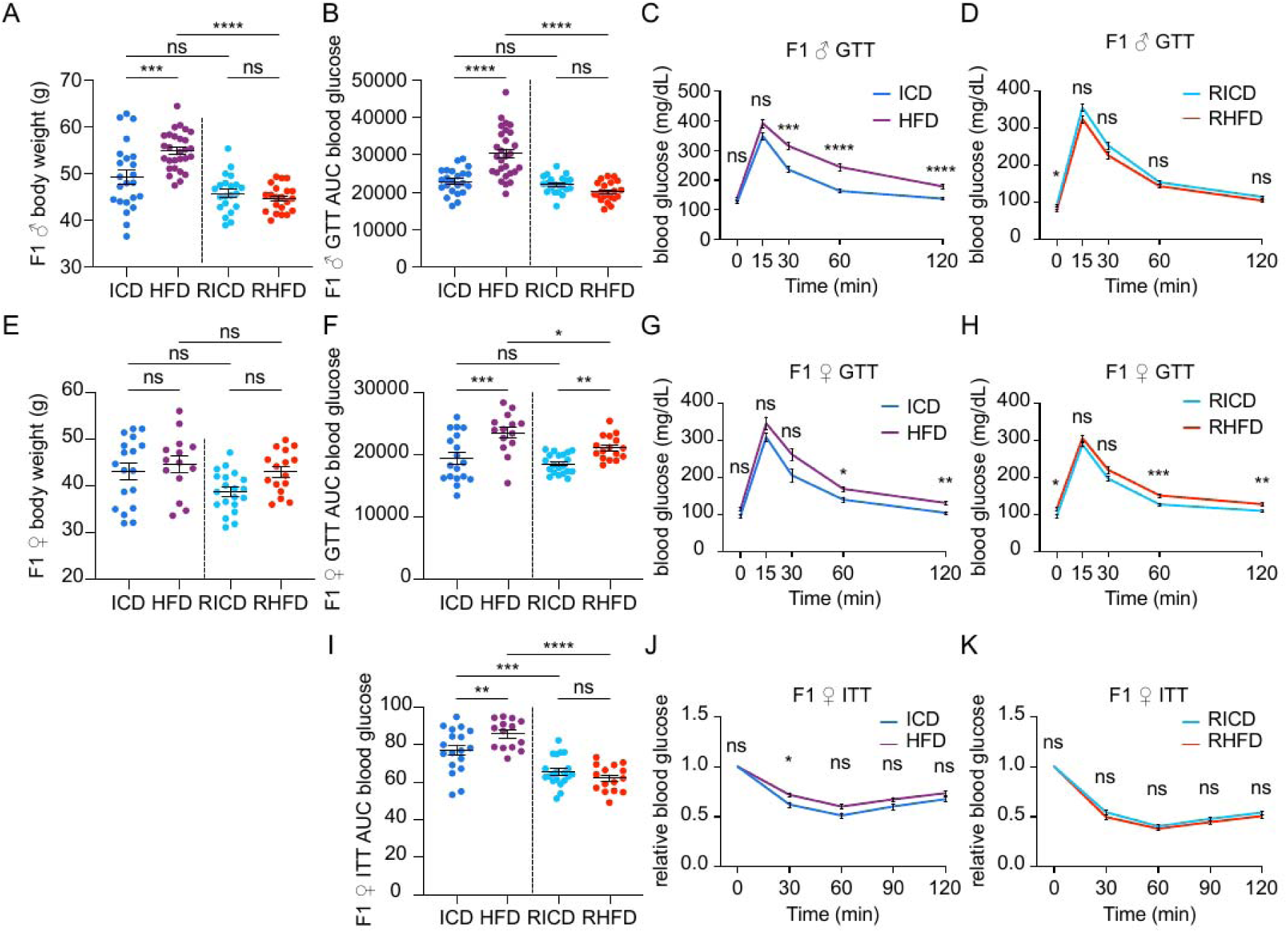
Intergenerational metabolic effects in F1 offspring before and after paternal reversal. (A and B) Body weight (A) and GTT AUC (B) in F1 males from ICD, HFD, RICD, and RHFD sires. (C and D) GTT time curves for F1 males. (E and F) Body weight (E) and GTT AUC (F) in F1 females. (G and H) GTT time curves for F1 females. (I) ITT AUC in F1 females. (J and K) ITT time curves for F1 females. F1 offspring from HFD sires showed sex-dependent metabolic impairment despite ICD rearing, whereas offspring from RHFD sires showed substantial normalization. Data are mean ± SEM. Statistical analyses accounted for offspring from multiple litters where applicable. *, **, ***, ****, and ns indicate p < 0.05, p < 0.01, p < 0.001, p < 0.0001, and not significant, respectively.

## Discussion

This study establishes a within-sire paternal induction-and-reversal model and uses it to show that intergenerational metabolic risk associated with paternal obesity is substantially reversible. The same HFD-exposed males that transmitted metabolic dysfunction to offspring before recovery subsequently sired offspring with improved metabolic outcomes after diet and exercise reversal. By using the same paternal lines before and after recovery, this design reduces confounding from paternal genetic background and founder-to-founder variation and directly links offspring phenotype to paternal metabolic state at conception. The RNA profiling data provide a molecular framework for this reversibility. HFD-Ex remodeled sperm mRNAs, lncRNAs, and sncRNAs, with sperm showing the strongest RNA response among the five tissues examined. Because mature sperm are transcriptionally inactive, these RNA changes most likely reflect altered germ-cell development, RNA processing, and RNA packaging during spermatogenesis. This interpretation is consistent with evidence that diet-induced heritable information is already present in testicular sperm and does not require epididymal transit.^28^ Reversal of paternal metabolic dysfunction was accompanied by broad restoration of sperm RNA profiles, suggesting that the testicular environment can be reset across subsequent spermatogenic cycles.

Different RNA classes showed different degrees of recovery. Sperm sncRNAs were almost completely restored after reversal, whereas mRNAs and lncRNAs showed partial residual differences. Although lncRNAs and sncRNAs exhibited a higher degree of recovery compared to mRNAs, this difference should be interpreted cautiously because these two types of noncoding RNAs are generally expressed at lower levels than mRNAs, which may reduce statistical power to detect persistent differential expression. Thus, the apparent recovery rates likely reflect both biological reversibility and RNA-class-specific differences in abundance and detection sensitivity. Somatic tissues also recovered broadly, in line with normalization of systemic metabolic indices. The residual differences in sperm mRNA and lncRNA after reversal may reflect a molecular record of prior HFD exposure rather than a continuing disease state. We therefore interpret these post-reversal differences cautiously: they may serve as biomarkers of exposure history, but their causal contribution to offspring phenotype remains to be tested.

The offspring data suggest that HFD-responsive sperm RNA changes corrected by reversal, rather than all residual post-reversal differences, are most closely associated with metabolic-risk transmission. F1 offspring conceived before reversal showed glucose intolerance and altered insulin sensitivity despite control-diet rearing, whereas offspring conceived after paternal recovery showed substantial metabolic improvement. This concordance between sperm sncRNA normalization and offspring rescue is consistent with a role for sncRNAs as mediators of the reversible signal.^21^ Functional testing will be required to define causality, including RNA injection, selective depletion or enrichment of candidate RNA classes, and comparison of sperm RNA fractions before and after reversal.

The sex dependence of the offspring response is also important. HFD-sired males exhibited increased body weight and impaired glucose tolerance, whereas HFD-sired females showed prominent glucose intolerance and altered insulin sensitivity with less consistent adiposity. After paternal reversal, male offspring were more fully normalized than female offspring. Such sexual dimorphism is common in paternal epigenetic inheritance and likely reflects sex-specific developmental, placental, endocrine, or metabolic responses to the same paternal signal.^33–35^ Future work should examine early embryos and placentas from male and female conceptuses to determine when these trajectories diverge.

These findings support a clear translational implication: paternal metabolic health before conception is a modifiable determinant of offspring metabolic risk. The use of outbred CD1 mice enhances the model’s relevance to genetically heterogeneous populations. In humans, spermatogenesis spans approximately 70 days, providing a plausible window during which diet, exercise, and metabolic interventions could alter the composition of sperm produced before conception. Because sperm DNA methylation has shown limited and inconsistent associations with paternal BMI in humans,^36^ sperm RNA should be prioritized in human preconception cohorts designed to evaluate paternal interventions.

In summary, paternal HFD-Ex induces metabolic dysfunction, remodels sperm RNA, and increases metabolic risk in offspring (Figure 6). Restoring paternal metabolic health before conception remodels the sperm RNA payload and substantially reduces intergenerational transmission of metabolic dysfunction (Figure 6). These findings support a model in which paternal epigenetic inheritance is not irreversibly fixed by prior exposure history but remains responsive to preconception metabolic recovery.

**Figure 6.**
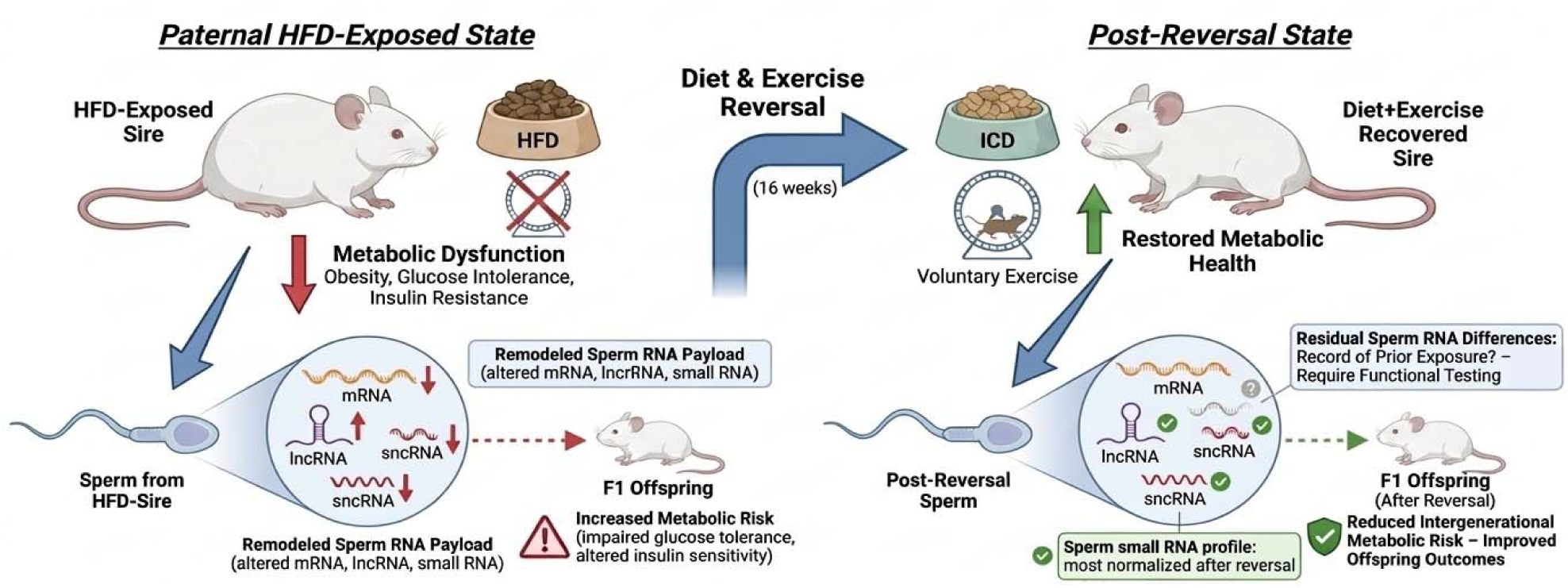
Schematic summary of the major findings of the present study. Paternal HFD-Ex induces obesity and remodels sperm mRNA, lncRNA, and sncRNA payloads, thereby increasing metabolic risk in offspring. Diet and exercise reversal restores paternal metabolic health, resets much of the sperm RNA landscape, and reduces intergenerational transmission of adverse metabolic traits. Residual sperm RNA differences after reversal may record prior exposure history, but their functional significance remains to be tested.

### Limitations of the study

First, although this study identifies sperm RNA remodeling as a molecular correlate of reversible intergenerational inheritance, it does not establish which RNA species are necessary or sufficient for transmission. RNA injection, ICSI using RNA-manipulated sperm, and selective depletion or enrichment of candidate RNA classes will be needed for causal testing. Second, the study was designed as an RNA-focused analysis and did not evaluate sperm DNA methylation, chromatin accessibility, or histone modifications. These layers may interact with sperm RNA and should be examined in future work. Third, multigenerational follow-up was not performed, so the persistence or further erasure of paternal exposure marks beyond F1 remains unknown. Fourth, natural mating does not exclude contributions from seminal plasma or paternal effects on the female reproductive tract. Although prior ICSI work supports a testicular sperm-borne mechanism,^28^ future studies should separate sperm-intrinsic and seminal-plasma contributions in the reversal setting.

## STAR Methods

### Key resources table

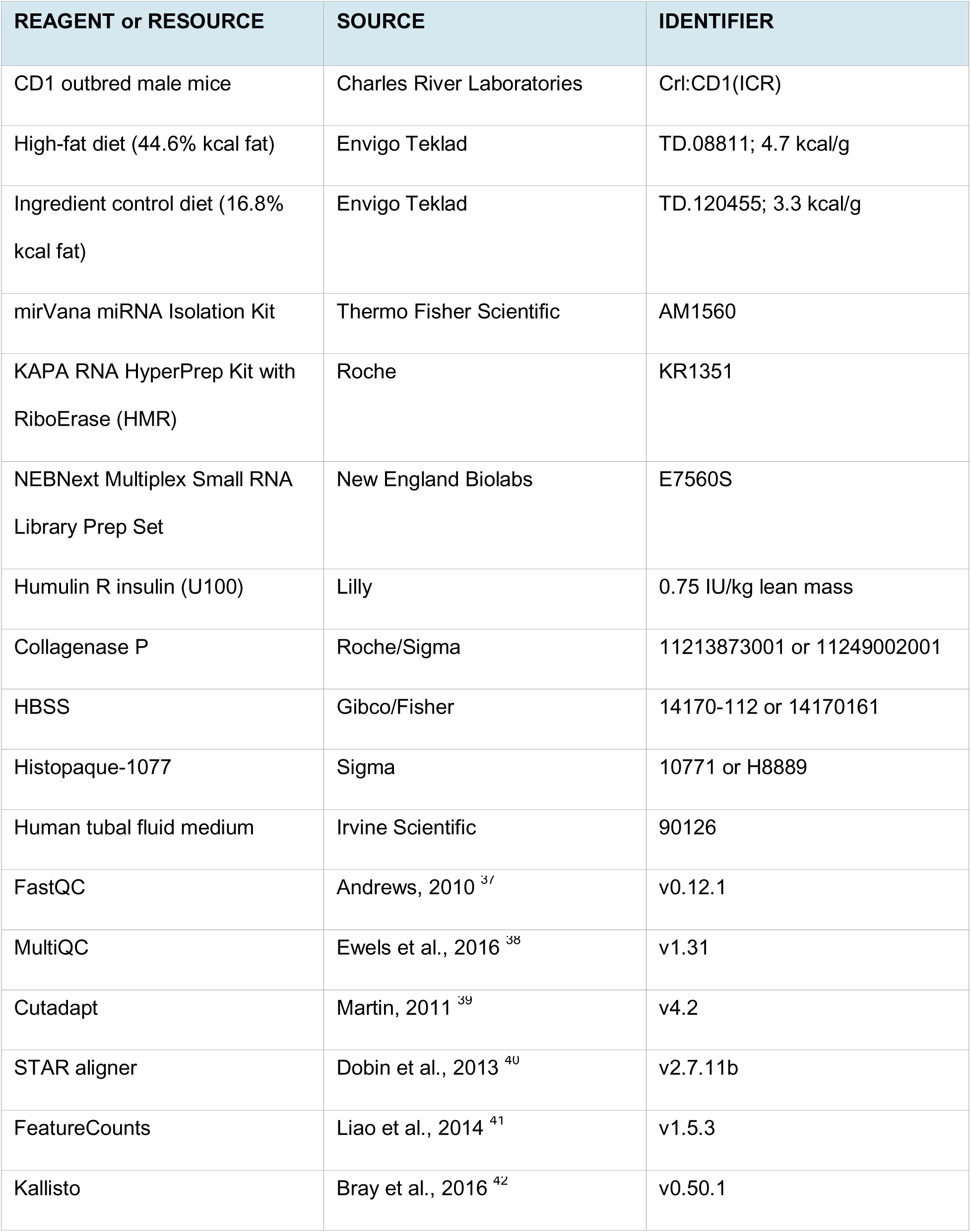
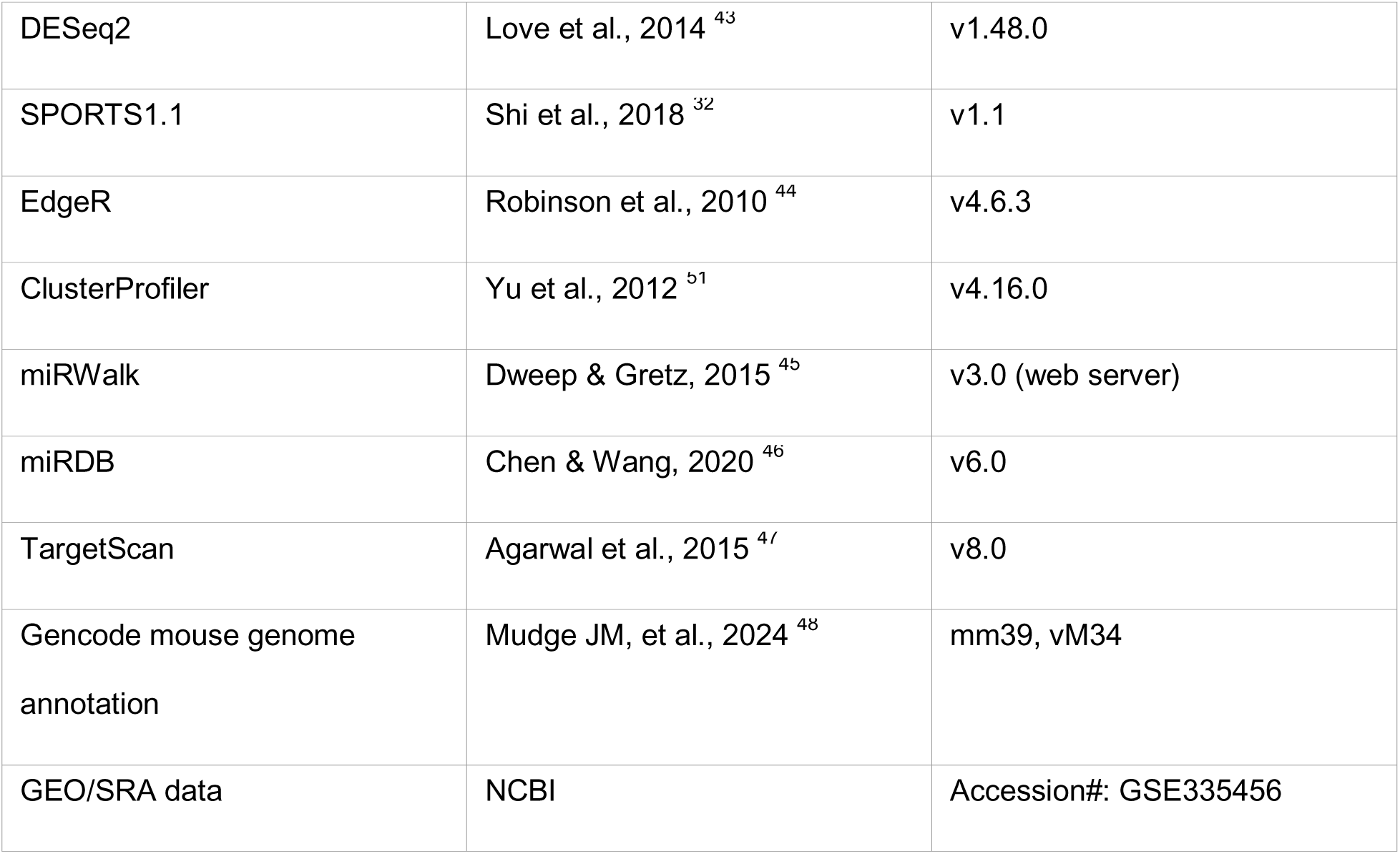

### Resource availability

#### Lead contact

Further information and requests for resources and reagents should be directed to and will be fulfilled by the Lead Contact, Wei Yan.

#### Materials availability

This study did not generate new unique mouse lines or plasmids. Biological materials will be made available upon reasonable request.

#### Data and code availability

RNA-seq and small RNA-seq datasets have been deposited in NCBI GEO/SRA under accession#: GSE335456. No custom software was generated. All analyses used established tools listed in the Key Resources Table.

### Experimental model and study participant details

#### Animals

Outbred CD1 F0 male mice (Charles River Laboratories, Crl:CD1(ICR)) were purchased at 3 weeks of age, acclimated and randomized at approximately 4 weeks of age. Males were housed individually with a running-wheel apparatus. All procedures were performed at C. W. Steers Biological Resources Center in accordance with IACUC guidelines of The Lundquist Institute for Biomedical Innovation at Harbor-UCLA Medical Center (Ref#050855). Animals were maintained under a 12 h light/12 h dark cycle at 22 ± 2°C with ad libitum access to water and assigned diets.

#### Diet and exercise intervention

Voluntary wheel running was performed as described^49^. Briefly, the basic protocol consists of allowing mice to run freely on the open surface of a slanted, saucer-shaped plastic wheel placed inside a standard rat cage. Rotations are transmitted via a USB hub so that the frequency and rotation rate can be captured by a software program for data storage and analysis over variable time periods. Mice are individually housed to ensure accurate recordings for each animal.

HFD and the ICD chow were purchased from Envigo (Table S8). HFD-Ex males received Envigo Teklad TD.08811 (44.6% kcal fat, 40.7% kcal carbohydrate, 14.7% kcal protein; 4.7 kcal/g) with the running wheels locked (-Ex) for 16 weeks. ICD+Ex males received Envigo Teklad TD.120455 (16.8% kcal fat, 60.9% kcal carbohydrate, 22.3% kcal protein; 3.3 kcal/g) with voluntary wheel running unlocked (+Ex) for 16 weeks. Body weight and running distance were recorded weekly. After initial phenotyping and first breeding, HFD-Ex males were switched to ICD+Ex for at least 16 weeks (RHFD), whereas controls remained on ICD+Ex (RICD). The F0 mice analyzed in this study are summarized in Table S7.

#### F1 offspring generation and maintenance

After phenotyping, F0 males were naturally mated (timed mating) with virgin CD1 females to generate F1 offspring before reversal (from ICD+Ex and HFD-Ex sires) and after reversal (from RICD and RHFD sires) (Table S7). The plugged females were separated to single cages the next morning and maintained on regular chow until parturition. F1 males and females were weaned at 3 weeks and maintained on regular chow without exercise until metabolic phenotyping at 4 and 10 months of age and tissue collection at 10 months of age (Tables S1 and S7). Offspring were derived from multiple sires and litters; sire-and litter-aware statistical models were used for F1 analyses where sample structure permitted. The F1 mice analyzed in this study are summarized in Tables S1 and S7.

## Method details

### DXA scanning, glucose tolerance, and insulin tolerance testing

Body composition was assessed by dual-energy X-ray absorptiometry (DXA) using a Hologic QDR 4500 Delphi whole-body DXA system (Hologic Inc., Bedford, MA, USA) before metabolic testing to determine fat mass and lean mass (Tables S1 and S7). For intraperitoneal glucose tolerance testing (GTT), mice were fasted for 6 h, from 9:00 a.m. to 3:00 p.m., and injected intraperitoneally with glucose at 2 g/kg lean mass. Blood glucose was measured from tail-vein blood using a OneTouch Ultra glucometer at 0, 15, 30, 60, and 120 min after glucose injection. For insulin tolerance testing (ITT), non-fasted mice were injected intraperitoneally with Humulin R insulin (U100) at 0.75 IU/kg lean mass. Blood glucose was measured from tail-vein blood at 0, 30, 60, 90, and 120 min after insulin injection. Area under the curve (AUC) was calculated using the trapezoidal method. Data are presented in Tables S1 and S7.

### Sperm collection and purification

Cauda epididymides were dissected into 1 mL human tubal fluid (HTF) medium in a 35-mm Petri dish. Small incisions were made at the proximal end using a 26G needle, and luminal contents were gently expressed with forceps. Samples were incubated at 37°C for 30 min to allow sperm release. Approximately 900 μL sperm-containing suspension was collected, and an additional 500 μL 1× phosphate-buffered saline (PBS) was used to recover remaining sperm while avoiding tissue debris. The combined suspension was centrifuged at 2,000–3,000 × g for 5 min. The pellet was resuspended in somatic cell lysis buffer (SCLB; 0.125 mL 10% SDS, 0.5 mL 0.5% Triton X-100, and 49.375 mL 1× PBS) and incubated on ice for approximately 1 min. Somatic cell lysis and sperm purity were confirmed by microscopy. Samples were centrifuged again at 2,000–3,000 × g for 5 min, the supernatant was discarded, and the sperm pellet was washed twice with 1× PBS. Final sperm pellets were either used immediately for RNA extraction or snap-frozen in liquid nitrogen and stored at −80°C.

### Sperm total RNA extraction

Mice used for transcriptomic analyses are listed in Table S7. Total RNA was extracted from purified sperm using the mirVana miRNA Isolation Kit (Thermo Fisher Scientific, AM1560) according to the manufacturer’s protocol with modifications. Briefly, 600 μL Lysis/Binding Buffer was added to each sperm pellet, and samples were homogenized until no visible clumps remained. One-tenth volume miRNA Homogenate Additive was added, mixed by inversion, and incubated on ice for 10 min. An equal volume of Acid-Phenol:Chloroform was then added, and samples were vortexed for 30–60 s and centrifuged at 15,000 × g for 20 min at 4°C. The aqueous phase was transferred to a fresh tube, mixed with 3 μL GlycoBlue and an equal volume of 2-propanol, and incubated at −20°C for at least 10 min. RNA was pelleted by centrifugation at 15,000 × g for 15 min at 4°C, washed twice with ice-cold 75% ethanol, and centrifuged at 10,000 × g for 5 min per wash. Pellets were air-dried and resuspended in 30 μL RNase-free water. RNA quality was assessed using an Agilent Bioanalyzer.

### Collection of pancreatic islets, skeletal muscle, liver, and brain

Pancreatic islets were isolated using an established collagenase perfusion method with minor modifications.^50^ Briefly, mice were euthanized, and the common bile duct was cannulated for intraductal perfusion of the pancreas with 0.5 mg/mL collagenase P (Roche, 11213873001) diluted in Hank’s balanced salt solution (HBSS; Gibco, 14170112). The inflated pancreas was dissected and incubated at 37°C for approximately 21 min to digest exocrine tissue. Digested pancreatic tissue was then washed and subjected to density-gradient separation using Histopaque-1077 (Sigma, H8889) by centrifugation at 900 × g for 15 min. Intact pancreatic islets were manually selected under a stereomicroscope to minimize exocrine contamination. Purified islets were washed in PBS, collected by gentle centrifugation, and either processed immediately for RNA extraction or snap-frozen in liquid nitrogen and stored at −80°C until use. Skeletal muscle, liver, and brain were collected from F0 males after metabolic phenotyping. Tissues were rapidly dissected, rinsed as appropriate to remove blood contamination, snap-frozen in liquid nitrogen, and stored at −80°C until nucleic acid extraction. Tissue collection was performed consistently across ICD+Ex, HFD-Ex, RICD, and RHFD groups to minimize technical variation (Table S7).

### Total RNA extraction from pancreatic islets and somatic tissues

Total RNA was extracted from isolated pancreatic islets or approximately 200 mg of brain, liver, or skeletal muscle using the mirVana miRNA Isolation Kit (Thermo Fisher Scientific, AM1560) with modifications for low-input samples. Samples were lysed in 600 μL Lysis/Binding Buffer, treated with miRNA Homogenate Additive, extracted with Acid-Phenol:Chloroform, and precipitated with GlycoBlue and 2-propanol at −20°C. RNA pellets were washed with 75% ethanol, air-dried, and resuspended in RNase-free water. RNA quality and size distribution were assessed using an Agilent Bioanalyzer.

### Library preparation and sequencing

For bulk RNA-seq, 100 ng total RNA from each sample was used for library construction using the KAPA RNA HyperPrep Kit with RiboErase (HMR) (Roche, KR1351), following the manufacturer’s protocol. Ribosomal RNA was depleted by DNA oligo hybridization and enzymatic digestion using RNase H and DNase. RNA was then fragmented and converted into first-and second-strand cDNA. Libraries were generated by A-tailing, ligation of Illumina-compatible adapters, and high-fidelity PCR amplification. Library concentration and fragment-size distribution were assessed before sequencing. Libraries were sequenced on an Illumina NovaSeq 6000 platform using paired-end 50 or 150-bp reads.

Small RNA-seq libraries were constructed from 100 ng total RNA using the NEBNext Multiplex Small RNA Library Prep Set for Illumina (New England Biolabs, E7560S), following the manufacturer’s recommendations. Briefly, 3′ and 5′ adapters were sequentially ligated to small RNAs, followed by reverse transcription and PCR amplification using barcoded primers. Final libraries were size-selected by PAGE gel excision or Pippin Prep to enrich fragments corresponding to 21–30 nt RNA inserts. Libraries were quality-controlled using an Agilent Bioanalyzer, quantified, and sequenced on an Illumina NovaSeq 6000 platform using single-end 50-bp reads.

### RNA-seq quality control

Raw sequencing read quality was assessed using FastQC,^37^ and quality-control metrics across all samples were summarized using MultiQC.^38^ Adapter sequences and low-quality bases were removed using Cutadapt ^39^ before alignment and downstream analysis.

### mRNA alignment, quantification, and normalization

Bulk RNA-seq reads were aligned to the GENCODE mm39 mouse reference genome (vM34) using STAR. ^40^ Exon-level read counts were generated using featureCounts ^41^ based on the GENCODE annotations. Raw count data were normalized using the relative log expression (RLE) method in DESeq2. ^43^ Protein-coding genes with at least 10 total counts across all samples were retained for downstream analysis.

### lncRNA alignment, quantification, and normalization

LncRNA expression was quantified from the same rRNA-depleted bulk RNA-seq libraries used for mRNA analysis. Transcript-level abundance estimates were generated using Kallisto ^42^ against the GENCODE mouse transcriptome reference corresponding to the mm39 genome and annotation version vM34. Kallisto-derived transcript-level estimated counts and abundances were imported into R using tximport and summarized to the gene level. Gene-level estimated counts were used to construct a DESeqDataSet using DESeqDataSetFromTximport. LncRNA genes were identified based on GENCODE gene biotype annotations by selecting genes annotated with the biotype term “lncRNA”. Low-abundance lncRNAs were removed before differential expression analysis using the same count-filtering strategy applied to protein-coding genes with at least 10 total counts across all samples.

### sncRNA alignment, annotation, and filtering

For sncRNA-seq, raw reads were aligned to the mm10 mouse genome using the SPORTS1.1 pipeline ^32^ with default settings and a maximum of one mismatch.

Reference annotations included miRNAs from miRBase v21,^51^ rRNAs from NCBI, tRNAs from GtRNAdb^52^, piRNAs from piRBase^53^ and piRNABank^54^, and other non-coding RNAs from Ensembl release 89 and Rfam 12.3.^55^ Count tables generated by SPORTS1.1 were annotated into sncRNA biotypes, including antisense RNAs, miRNAs, mitochondrial rRNA-derived small RNAs (mt-rsRNAs), mitochondrial tRNA-derived small RNAs (mt-tsRNAs), tRNA-derived small RNAs (tsRNAs), rRNA-derived small RNAs (rsRNAs), piRNAs, scaRNAs, snRNAs, snoRNAs, and YRNAs.^32^ Low-abundance features were removed using the filterByExpr() function in edgeR (v4.2.2)^44^ with default parameters while accounting for experimental group structure.

### Differential expression analysis

Differential expression of protein-coding genes was performed using DESeq2 (v1.48.0).^43^ Genes were considered significantly differentially expressed if they passed Benjamini–Hochberg multiple-testing correction with an adjusted p-value < 0.05.^56^ A log2 fold-change threshold of |0.3| was incorporated directly into the statistical test using the lfcThreshold parameter in the DESeq2 results() function. Differential expression of lncRNAs was assessed using DESeq2 with the same model structure, filtering criteria, multiple-testing correction, and log2 fold-change threshold as for protein-coding genes.

Differential expression of sncRNA fragments was assessed using edgeR’s glmTreat framework,^44^ which tests whether expression changes exceed a predefined biological effect size. sncRNAs were considered differentially expressed when they showed |log2 fold change| ≥ 0.3 and passed Benjamini–Hochberg correction with FDR < 0.05.

### miRNA target prediction

Predicted target genes of differentially expressed miRNAs were identified using the miRWalk web server.^45^ Target prediction was restricted to 3′ untranslated regions (3′ UTRs), the canonical sites of miRNA–mRNA interaction. Only target genes predicted by miRWalk, miRDB,^46^ and TargetScan,^47^ and with a miRWalk binding probability greater than 95%, were retained for downstream functional analysis.

### Functional enrichment analysis

Gene Ontology (GO) enrichment analysis was performed using the PANTHER Classification System overrepresentation test through the Gene Ontology Consortium resource.^42,57,58^ GO Biological Process annotations for *Mus musculus* were used. Statistical significance was assessed using Fisher’s exact test with Bonferroni correction for multiple testing. GO annotations correspond to the Gene Ontology release accessed on January 15, 2026.

### Quantification and statistical analysis

Data are presented as mean ± SEM unless otherwise stated. F0 comparisons were performed using two-tailed Student’s t tests or one-way ANOVA followed by Tukey post hoc correction, as appropriate. For F1 offspring, analyses accounted for relatedness among offspring from the same sire and litter. Linear mixed-effects models included treatment group as a fixed effect and sire and litter as random effects when the data structure supported this approach; otherwise, litter means were analyzed. Male and female offspring were analyzed separately unless stated otherwise. For GTT and ITT time courses, treatment, time, and treatment-by-time interaction were modeled as fixed effects, with repeated measures accounted for by offspring ID. RNA-seq and sncRNA-seq significance was defined as Benjamini-Hochberg-adjusted p < 0.05. Figure symbols denote: ns, not significant; *p < 0.05; **p < 0.01; ***p < 0.001; ****p < 0.0001.

## Supplemental Information

## Supplemental Figures

Figure S1. Validation of representative HFD-responsive sperm mRNAs.

Figure S2. sncRNA diversity across sperm and metabolic tissues.

Figure S3. Composition of differentially expressed sperm sncRNAs before and after paternal metabolic reversal.

Figure S4. Tissue distribution of differentially expressed sncRNAs before and after paternal metabolic reversal.

Figure S5. Correlation analysis of sperm sncRNA profiles across paternal treatment groups.

## Supplemental Datasets

Table S1. F0 and F1 metabolic profiling data.

Table S2. Differentially expressed mRNAs in F0 sperm and metabolic tissues.

Table S3. GO term enrichment analyses on mRNA DEGs.

Table S4. Differentially expressed lncRNAs in F0 sperm and metabolic tissues.

Table S5. Differentially expressed sncRNAs F0 sperm and metabolic tissues.

Table S6. miRNA target prediction and GO enrichment results.

Table S7. F0 and F1 mice analyzed in this study.

Table S8. Macronutrients of the ingredient control diet (ICD) and high-fat diet (HFD) purchased from Envigo

## Acknowledgments

We thank the staff of C. W. Steers Biological Resources Center at The Lundquist Institute for animal care support. This work was supported by grants from the NIH/NICHD (HD098593 to W.Y.), NIH/NCATS UCLA CTSI (UL1TR001881-01 to Z.W. and W.Y.), NIH/NIDDK (R01DK136888 to E.Y.), California Institute for Regenerative Medicine (EDUC4-12837 to Z.W. and W.Y.), and Washington State University startup funds (PG00023119 to W.Y.).

## Author Contributions

Conceptualization, W.Y., H. R., R.B., C.W., R.S.S., J.R.M.; methodology and investigation, S.C., H.Z., R.W., H.M., F.C., J.C., E.Y., H.R., R.B., Y.L., C.W., R.S.S., L.C. and W.Y.; formal analysis, R.D.M.M., R.W., Z.W.; visualization, R.D.M.M. and Z.W.; writing - original draft, W.Y.; writing - review and editing, all authors; funding acquisition, W.Y.; supervision, W.Y. Author contributions follow CRediT taxonomy.

