## Supplementary material for "Paternal Metabolic Reversal Remodels Sperm RNA Profiles and Ameliorates Intergenerational Metabolic Disorder in Mice": Figures S1-S5

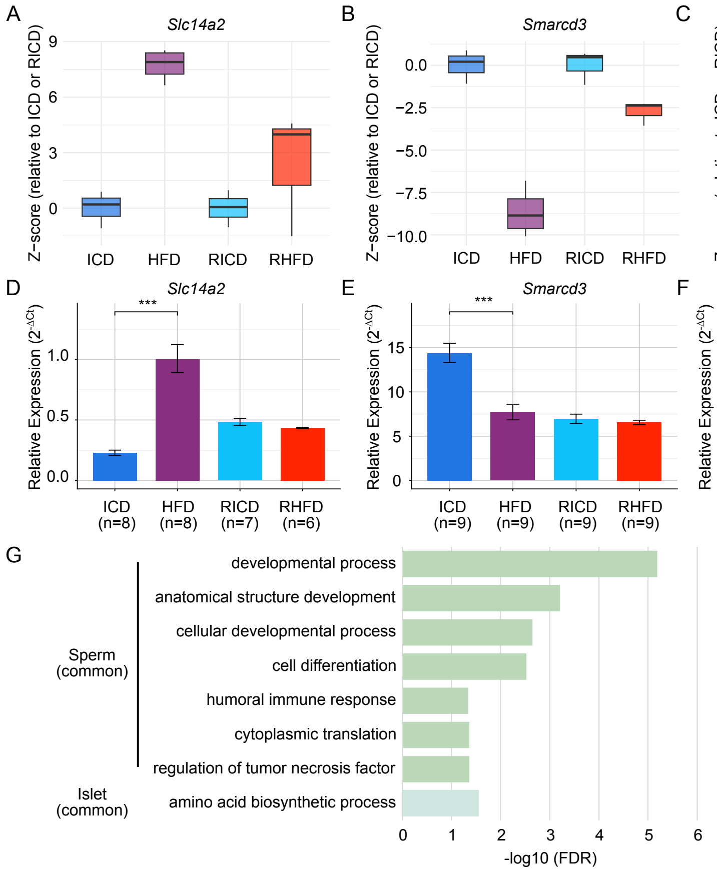

A
Slc14a2
Z-score (relative to ICD or RICD)
ICD
HFD
RICD
RHFD
B
Smarcd3
C
Srek1
D
Slc14a2
Relative Expression (2^-ΔCt)
ICD (n=8)
HFD (n=8)
RICD (n=7)
RHFD (n=6)
E
Smarcd3
ICD (n=9)
HFD (n=9)
RICD (n=9)
RHFD (n=9)
F
Srek1
ICD (n=4)
HFD (n=2)
RICD (n=9)
RHFD (n=4)
G
Sperm (common)
Islet (common)
developmental process
anatomical structure development
cellular developmental process
cell differentiation
humoral immune response
cytoplasmic translation
regulation of tumor necrosis factor
amino acid biosynthetic process
-log10 (FDR)

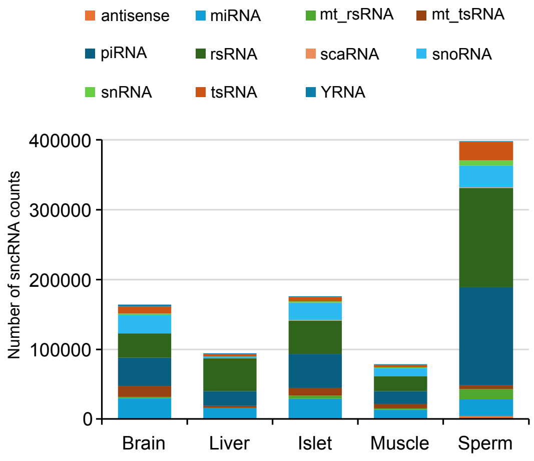
antisense
miRNA
mt_rsRNA
mt_tsRNA
piRNA
rsRNA
scaRNA
snoRNA
snRNA
tsRNA
YRNA
400000
300000
200000
100000
0
Number of sncRNA counts
Brain
Liver
Islet
Muscle
Sperm

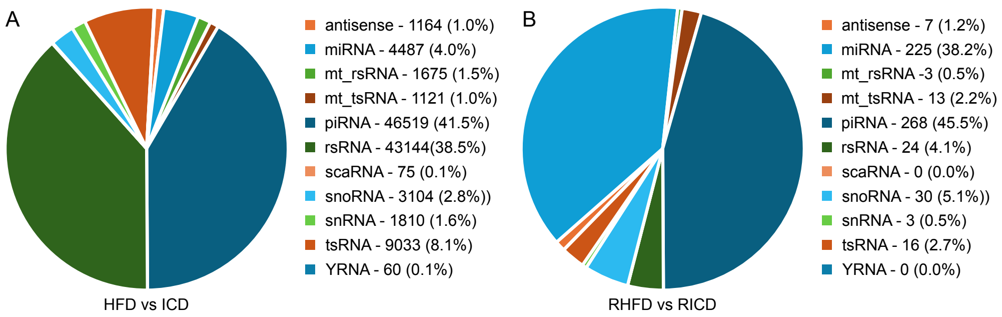
A
antisense - 1164 (1.0%)
miRNA - 4487 (4.0%)
mt_rsRNA - 1675 (1.5%)
mt_tsRNA - 1121 (1.0%)
piRNA - 46519 (41.5%)
rsRNA - 43144(38.5%)
scaRNA - 75 (0.1%)
snoRNA - 3104 (2.8%))
snRNA - 1810 (1.6%)
tsRNA - 9033 (8.1%)
YRNA - 60 (0.1%)
HFD vs ICD
B
antisense - 7 (1.2%)
miRNA - 225 (38.2%)
mt_rsRNA -3 (0.5%)
mt_tsRNA - 13 (2.2%)
piRNA - 268 (45.5%)
rsRNA - 24 (4.1%)
scaRNA - 0 (0.0%)
snoRNA - 30 (5.1%))
snRNA - 3 (0.5%)
tsRNA - 16 (2.7%)
YRNA - 0 (0.0%)
RHFD vs RICD

A

| HFD vs ICD | Brain | Islet | Liver | Muscle | Sperm |
|---|---|---|---|---|---|
| antisense | 0 | 0 | 0 | 0 | 1164 |
| miRNA | 0 | 1 | 0 | 15 | 4487 |
| mt_rsRNA | 0 | 0 | 0 | 1 | 1675 |
| mt_tsRNA | 0 | 0 | 0 | 24 | 1121 |
| piRNA | 0 | 21 | 0 | 9 | 46519 |
| rsRNA | 0 | 1 | 0 | 73 | 43144 |
| scaRNA | 0 | 0 | 0 | 2 | 75 |
| snoRNA | 0 | 0 | 0 | 30 | 3104 |
| snRNA | 0 | 0 | 0 | 15 | 1810 |
| tsRNA | 0 | 0 | 0 | 17 | 9033 |
| YRNA | 0 | 0 | 0 | 0 | 60 |
| TOTAL | 0 | 23 | 0 | 186 | 112192 |

B

| RHFD vs RICD | Brain | Islet | Liver | Muscle | Sperm |
|---|---|---|---|---|---|
| antisense | 0 | 0 | 0 | 0 | 7 |
| miRNA | 1 | 0 | 0 | 0 | 225 |
| mt_rsRNA | 0 | 0 | 0 | 0 | 3 |
| mt_tsRNA | 0 | 0 | 0 | 0 | 13 |
| piRNA | 0 | 0 | 0 | 0 | 268 |
| rsRNA | 0 | 0 | 0 | 0 | 24 |
| scaRNA | 0 | 0 | 0 | 0 | 0 |
| snoRNA | 0 | 0 | 0 | 0 | 30 |
| snRNA | 0 | 0 | 0 | 0 | 3 |
| tsRNA | 0 | 0 | 0 | 0 | 16 |
| YRNA | 0 | 0 | 0 | 0 | 0 |
| TOTAL | 1 | 0 | 0 | 0 | 589 |

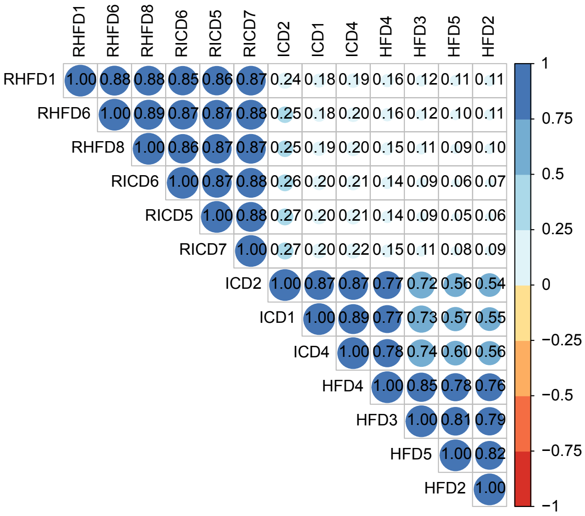
RHFD1 RHFD6 RHFD8 RICD6 RICD5 RICD7 ICD2 ICD1 ICD4 HFD4 HFD3 HFD5 HFD2
RHFD1 1.00 0.88 0.88 0.85 0.86 0.87 0.24 0.18 0.19 0.16 0.12 0.11 0.11
RHFD6 1.00 0.89 0.87 0.87 0.88 0.25 0.18 0.20 0.16 0.12 0.10 0.11
RHFD8 1.00 0.86 0.87 0.87 0.25 0.19 0.20 0.15 0.11 0.09 0.10
RICD6 1.00 0.87 0.88 0.26 0.20 0.21 0.14 0.09 0.06 0.07
RICD5 1.00 0.88 0.27 0.20 0.21 0.14 0.09 0.05 0.06
RICD7 1.00 0.27 0.20 0.22 0.15 0.11 0.08 0.09
ICD2 1.00 0.87 0.87 0.77 0.72 0.56 0.54
ICD1 1.00 0.89 0.77 0.73 0.57 0.55
ICD4 1.00 0.78 0.74 0.60 0.56
HFD4 1.00 0.85 0.78 0.76
HFD3 1.00 0.81 0.79
HFD5 1.00 0.82
HFD2 1.00
1
0.75
0.5
0.25
0
−0.25
−0.5
−0.75
−1
